# Predicting the structures of cyclic peptides containing unnatural amino acids by HighFold2

**DOI:** 10.1101/2025.01.16.633493

**Authors:** Cheng Zhu, Sen Cao, Tianfeng Shang, Jingjing Guo, An Su, Chengxi Li, Hongliang Duan

## Abstract

Cyclic peptides containing unnatural amino acids possess many excellent properties and have become promising candidates in drug discovery. Therefore, accurately predicting the three-dimensional structures of cyclic peptides containing unnatural residues will significantly advance the development of cyclic peptide-based therapeutics. Although deep learning-based structural prediction models have made tremendous progress, these models still cannot predict the structures of cyclic peptides containing unnatural amino acids. To address this gap, we introduce a novel model, HighFold2, built upon the AlphaFold-Multimer framework. HighFold2 first extends the pre-defined rigid groups and their initial atomic coordinates from natural amino acids to unnatural amino acids, thus enabling structural prediction for these residues. Then, it incorporates an additional neural network to characterize the atom-level features of peptides, allowing for multi-scale modeling of peptide molecules while enabling the distinction between various unnatural amino acids. Besides, HighFold2 constructs a relative position encoding matrix for cyclic peptides based on different cyclization constraints. Except for training using spatial structures with unnatural amino acids, HighFold2 also parameterizes the unnatural amino acids to relax the predicted structure by energy minimization for clash elimination. Extensive empirical experiments demonstrate that HighFold2 can accurately predict the three-dimensional structures of cyclic peptide monomers containing unnatural amino acids and their complexes with proteins, with the median RMSD for Cα reaching 1.891 Å. All these results indicate the effectiveness of HighFold2, representing a significant advancement in cyclic peptide-based drug discovery.

## 1. Introduction

Peptides are compounds composed of amino acids linked by peptide bonds^1^, with molecular weights between small molecules and macromolecules. Renowned for their ability to selectively modulate diverse protein-protein interactions, peptides are pivotal in drug development^2,3^. Peptide-based therapeutics offer unique advantages: compared to small-molecule drugs, they exhibit greater specificity and reduced toxicity; relative to protein-based drugs, they possess diminished immunogenicity and lower production costs^4^. These attributes position peptides as compelling candidates for treating a broad spectrum of diseases^5^. Nonetheless, linear peptides are susceptible to being hydrolyzed by proteases^6^. Cyclization, coupled with the incorporation of unnatural amino acids, can markedly enhance their stability, binding affinity, specificity, and membrane permeability, thus enabling them to target intracellular proteins effectively^7,8^. Given these distinctive benefits of cyclic peptides with unnatural amino acids, accurate prediction of their monomeric and complex spatial structures has become increasingly critical.

The rapid evolution of artificial intelligence algorithms has revolutionized structural prediction models. Tools such as AlphaFold2^9^, AlphaFold-Multimer^10^, RoseTTAFold^11^, and ESMFold^12^ have achieved remarkable accuracy in predicting the structures of protein monomers and protein-protein complexes. Building upon these milestones, subsequent advancements have extended predictive capabilities to encompass protein-nucleic acid^13^, protein-peptide^14^, protein-small molecule^15^, and protein-cyclic peptide complexes^16^. Recently, models like RoseTTAFold All-Atom^17^ and AlphaFold3^18^ have further broadened the horizon of deep learning-based spatial structure prediction to the joint structure of complexes involving proteins, nucleic acids, small molecules, ions, and modified residues. However, despite these advancements, current methodologies remain limited. They either predict structures for cyclic peptides containing natural amino acids or linear peptides with unnatural amino acids, none specifically address the structural prediction of cyclic peptides incorporating unnatural amino acids and their associated complexes.

In this study, we leveraged AlphaFold-Multimer to achieve accurate spatial structure predictions for cyclic peptides with unnatural amino acids and their complexes. Considering peptides have certain small molecule properties, we integrated a neural network that characterizes peptide atomic properties into AlphaFold-Multimer, enabling multi-scale modeling of peptide molecules and distinguishing different unnatural residues. Furthermore, we extended the rigid groups of natural amino acids to encompass unnatural amino acids. These enhancements allowed us to fine-tune the combined model using three-dimensional structural data containing unnatural amino acids. While this model could be directly trained on cyclic peptides with unnatural amino acids, the scarcity of such structures in the Protein Data Bank (PDB)^19,20^ necessitated an alternative approach. So, we use the zero-shot learning method, first fine-tuning the model using linear peptides with unnatural amino acids and then modifying the relative position encoding matrix to enable accurate cyclic peptides and their complexes predictions.

The prediction results demonstrate that HighFold2 can accurately predict the spatial structures of cyclic peptides containing unnatural amino acids and their complexes. Compared to the native structure, the median RMSD (root-mean-square deviation) between the Cα atoms in the predicted and native structures is 1.891 Å, while the median RMSD for all atoms in the structure reaches 2.872 Å, and the median RMSD between atoms in the unnatural amino acids is 2.579 Å. Subsequent ablation experiments further demonstrated the efficacy of the individual components of the model in predicting cyclic peptide structures with unnatural amino acids. Prediction results also showed that HighFold2 performed well on the test set of linear peptides containing unnatural amino acids. Additionally, we parameterized the unnatural amino acids and performed energy minimization to relax the predicted structures, thereby eliminating potential spatial clashes.

In summary, we have developed a method termed HighFold2 capable of accurately predicting the structures of cyclic peptides containing unnatural amino acids and their complexes. This method involves training a deep learning model based on AlphaFold-Multimer using linear peptide structures with unnatural amino acids, then modifying the model’s relative position encoding matrix, enabling it to predict the cyclic peptide structures successfully. Then, relaxation is performed to refine the spatial structure further. We believe that this method will serve as a powerful tool for the development of cyclic peptide-based therapeutics.

## 2. Methods and Materials

### 2.1 Datasets

In this study, we utilized two datasets: linear peptides with unnatural amino acids and cyclic peptides with unnatural amino acids. The linear peptide dataset with unnatural amino acids was used to train the model, while the cyclic peptide dataset with unnatural amino acids was used exclusively for testing the model.

The linear peptide data containing unnatural amino acids were sourced from the ModPep dataset^21^, which includes 501 samples of linear peptides with unnatural amino acid residues. Most of these are complexes formed from linear peptides with unnatural residues and proteins. Using the provided PDB IDs, we downloaded the corresponding crystal structures from the PDB database, removed duplicate entries, and ultimately obtained 419 unique three-dimensional structures. We removed solvents, hydrogen atoms, and heteroatoms from the crystal structures to facilitate subsequent model training and testing and retained only each structure’s first state.

To minimize redundancy in the crystal structures and reduce computational demands during training, we applied the following rules to remove certain chains from the spatial structure: chains with fewer than 50 amino acid residues were classified as peptide chains, while others were defined as protein chains. If a structure contained at least one protein chain, we aligned the sequences of all peptide chains. For peptide chains with identical or partially overlapping sequences separated by a distance greater than 5 Å, only the longest peptide chain was retained, and the others were discarded. Additionally, if at least two protein chains remained in this structure, we retained only the protein chain that contacts most with the peptide chain.

Then, we analyzed the types of unnatural amino acids in the crystal structures. Any structure containing the unnatural amino acid that was only found once was excluded. Samples containing nucleic acids or acetyl groups were also removed. After this screening, 382 samples remained, representing 23 distinct unnatural amino acid residues.

Besides, cyclic peptides with unnatural amino acids were sourced from the cPEPmatch webserver^22^. To effectively evaluate the model’s accuracy on cyclic peptide data, we filtered the dataset to retain samples that only contained the same 23 types of unnatural amino acid residues as in the linear peptide dataset. We excluded samples with other unnatural amino acids and limited the cyclization forms to head-to-tail cyclization or disulfide bridge formation. Additionally, we also searched the PDB database to identify some structures that met these criteria. After downloading the corresponding crystal structures from the PDB, we removed solvent molecules, hydrogen atoms, and heteroatoms, retaining only the first state of each structure. We also excluded samples containing nucleic acids and duplicate entries. In the end, we obtained a total of 34 samples of cyclic peptides containing unnatural amino acids.

### 2.2 The architecture

HighFold2 is based on AlphaFold-Multimer and fine-tuned with its parameters. To enable the prediction of the three-dimensional structures of peptide monomers containing unnatural amino acids and their complexes, we introduced two major modifications to the AlphaFold-Multimer architecture: the integration of a neural network module to extract atomic-scale features of peptide molecules and the extension of AlphaFold-Multimer’s rigid groups to include unnatural amino acids.

#### 2.2.1 Atomic-scale feature extraction

AlphaFold-Multimer originally extracts features of peptides at the residue level, omitting atomic-scale information. To address this limitation and distinguish various unnatural residues, we incorporated an additional neural network module to capture atomic-scale representations of peptides. These representations were then combined with the residue-level features from AlphaFold-Multimer to achieve multi-scale modeling, as shown in Fig. 1B. The atomic-scale features were derived from atom element and bond information, which was extracted by converting chains with fewer than 50 amino acid residues in crystal structures from FASTA^23^ to SMILES^24^ using RDKit.

Atom element features were encoded using one-hot encoding. To reduce the sparsity of the encoding matrix, we separately encoded common atoms (B, C, F, I, N, O, P, S, Br, Cl), while rare atoms were assigned a shared encoding. The resulting one-hot matrix was processed using a multi-head attention mechanism (MHA)^25^ defined as follows:

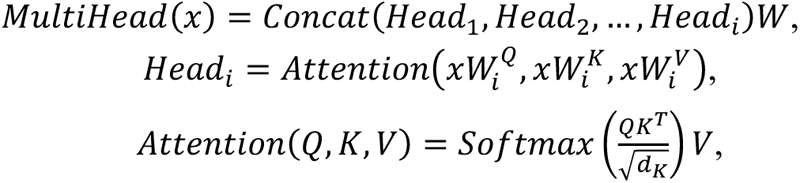

where, x represents the matrix of encoded atom element features with eight heads in total, Q, K, and V denote query, key, and value matrices, and d_K_ is the dimension of the key matrix. All Ws represent learnable parameters. Then, the multi-head attention outputs were passed through a multi-layer perceptron (MLP)^26^ to transform their dimensions to 21 (corresponding to the 20 natural amino acids and one unknown amino acid). The i-th layer of MLP is defined as:

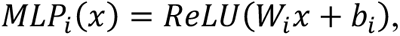

where x denotes the input a tom element feature embeddings of the i-th MLP layer. W_i_ and b_i_ are the learnable weights and biases.

To integrate atomic-scale features into AlphaFold-Multimer, we applied an attention-based pooling mechanism^27^, mapping atomic-level features to residue-level representations:

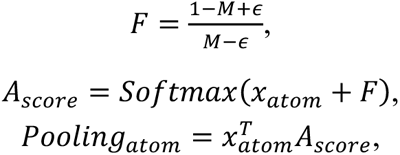

where M is the mask matrix (N × N_r_, where N is the number of atoms, and N_r_ is the number of residues), indicating the residue to which each atom belongs, ɛ is a small positive value to prevent division by zero, and x_atom_ represents atomic element embeddings after the MLP layers. The resulting Pooling_atom_ (N_r_ × 21) was added to the AlphaFold-Multimer’s one-hot matrix of peptide residue type and then input into the Evoformer module to generate the single representation. To keep it concise, we omit the dimensional extension and reduction operations of variables in the formula.

Bond information between atoms was also encoded using the one-hot approach, covering five bond types (single, double, triple, aromatic, and ionic bond). The resulting matrix (N × N × 5) was transformed via a linear layer to reduce its dimensionality to N × N × 1. Then, using a pooling mechanism like that described above, the bond information was mapped to residue-level representations and added to AlphaFold-Multimer’s pair activations matrix of peptide to generate the pair representation. These modifications enabled the model to effectively capture and integrate atomic-scale details into the residue-level framework of AlphaFold-Multimer, facilitating the prediction of three-dimensional structures for peptides and their complexes, including those with unnatural amino acids.

#### 2.2.2 Extend residue rigid groups to unnatural amino acids

The structure module in AlphaFold-Multimer leverages the single representation and pair representation derived from the Evoformer module to predict the torsion angles of each amino acid residue. Then, these predicted torsion angles, along with pre-defined rigid groups and atomic coordinates, facilitate the construction of all atomic coordinates in the spatial structure, simplifying the prediction complexity. We implemented a similar approach by defining rigid groups and initializing atomic coordinates for unnatural amino acids to extend the model to these residues, as shown in Fig. 1C.

Each atom within an amino acid residue can be categorized into one of five rigid groups based on its dependence on specific torsion angles. These groups include the backbone rigid group, which contains the backbone atoms Cα, Cβ, C, and N, while the ω and ϕ rigid groups, associated with the hydrogen atoms on Cα and the amino group, respectively, are excluded since our model does not predict hydrogen atoms. The ψ rigid group encompasses the oxygen atom in the carboxyl group, and the χ rigid group includes all side chain atoms. Specific details of the rigid group assignments for unnatural amino acids are documented in Table S1.

After defining the rigid groups, their initial coordinates are determined based on the crystallographic structures of unnatural amino acids. For the backbone rigid group, the coordinate system is centered at Cα, with C positioned along the positive x-axis, N in the xy-plane, and Cβ calculated relative to these references. The ψ rigid group is initialized with C as the origin, Cα on the negative x-axis, and the nitrogen of the subsequent residue within the xy-plane, allowing the position of the oxygen atom to be established. The side chain χ rigid group which are organized into four subgroups dependent on different torsion. Despite some amino acids having more than four torsion angles, smaller angles with negligible effects are disregarded. Within each subgroup, the third atom is set as the origin, the second atom is aligned to the negative x-axis, and the first atom lies in the xy-plane. The relative position of the fourth atom is rotated into the xy-plane using a rotation matrix Rx, as follows:

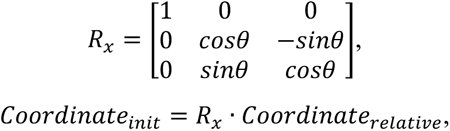

where θ represents the rotation angle between the relative position of the fourth atom and the xy-plane when it rotates around the x-axis.

#### 2.2.3 Modify relative position encoding for cyclization

To represent cyclic position information, we construct a cyclic position matrix by calculating the shortest distance for any two residues, as shown in Fig. 1D. For cyclic peptides with the head-to-tail constraint, the cyclic position can be formulated as:

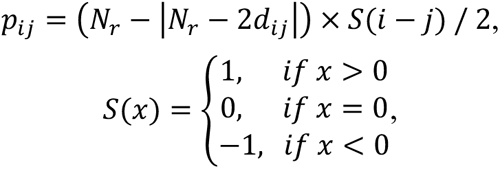

where *p_ij_* denotes the relative position between residue i and residue j in the cyclic position matrix, *d_ij_* denotes the relative distance between residue i and residue j, *N_r_* denotes the number of residues (including the unnatural residues), |·| denotes the symbol of absolute value.

For the scenario of cyclization with disulfide bridge constraints, the cyclic position matrix becomes more complicated due to the existence of disulfide bridges. As illustrated in HighFold^16^, the relative position between any two residues is calculated as the shortest distance between them in the residual topological graph based on the classic Floyd-Warshall algorithm. The signal assignment is conducted according to the directionality of amide bonds in the practical position offset. For brevity, we do not elaborate further here, and please refer to the HighFold model for details.

### 2.3 Training

HighFold2 is trained based on the dataset of linear peptides, dividing it into training, validation, and test sets at a ratio of 7.5:1.5:1. Each unnatural amino acid was ensured to appear at least once in both the validation and test sets. Features required for training and validation were generated using ColabFold^28^, while template features were excluded to prevent data leakage. Due to the limitation of computational resources, we cropped the number of amino acid residues in each training sample to 220 while the validation set remained uncropped. To mitigate overfitting, early stopping was implemented based on the loss of the validation set. We fine-tuned all five sets of parameters from AlphaFold-Multimer, allowing us to generate five predicted structures during testing, like the procedure of AlphaFold-Multimer.

### 2.4 Relaxation

#### 2.4.1 Parameterization of unnatural amino acid residues

To be compatible with Amber force fields^29^, force field parameters of unnatural amino acid residues were generated by following its official tutorial. In brief, each unnatural amino acid residue was capped with N-methyl and acetyl groups in the form of ACE-XXX-NME in GaussView. Structure optimization was then performed with Gaussian 16 by using density functional theory (DFT)^30^ at the B3LYP/6-31G(d) level, followed by a calculation at the level of HF/6-31G(d) on the optimized structure to reproduce the electrostatic potential^31^. Subsequently, the atomic partial charges of the unnatural amino acid residue were generated using the restricted electrostatic potential (RESP) method with Antechamber, during this process the charge of the capping groups was fixed. The penalty function of RESP is formulated as follows:

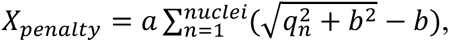

where α defines the asymptotic limit of the penalty, *q_n_* represents the partial charge of the *n*-th atom, and b is the width parameter.

The charge sets of the ACE and NME groups are identical to the Amber ff14SB force fields^32^. Afterward, the capping groups were removed to generate the force field parameters of the unnatural amino acid residue. Finally, the .prepin and .frcmod files were generated based on the General Amber Force Field (GAFF)^33^.

#### 2.4.2 Energy minimization

The PDB file is loaded in AmberTools^34^, and the structural topology and coordinate files were created by combining the Amber ff14SB, GAFF, and the generated unnatural amino acid residue parameters. For cyclic peptides, the ‘bond’ command is used to establish connections between the pair of atoms; for complex systems, ParmEd^35^ is employed to specify the chain names for each peptide chain. The generated topology and coordinate files are then loaded into OpenMM^36^ for an iterative restrained energy minimization procedure. Harmonic restraints with a spring constant of 10 kcal/mol·Å² are applied to heavy atoms to maintain proximity to the input structure. Violations are resolved by identifying residues with clashes, removing their restraints, and re-minimizing iteratively until all issues are addressed. The process is performed with OpenMM’s default tolerance of 2.39 kcal/mol and no step limit, ensuring structural integrity and proper hydrogen placement.

### 2.5 Metrics

#### 2.5.1 RMSD calculation

We evaluated the model’s predictive performance by calculating the root-mean-square deviation (RMSD) between the predicted and actual structures. Specifically, the pocket-aligned peptide RMSD_all-atom_, pocket-aligned peptide RMSD_Cα_, and pocket-aligned peptide RMSD_unAA_ were used to assess the structural accuracy of complexes containing unnatural amino acids, while RMSD_all-atom_, RMSD_Cα_, and RMSD_unAA_ were employed for evaluating the structures of peptide monomers with unnatural amino acids.

The pocket-aligned peptide RMSD_all-atom_ and pocket-aligned peptide RMSD_unAA_ are defined as follows: for each crystal structure, chains with fewer than 25 residues are identified as peptide chains, and the remaining chains are defined as protein chains. Backbone atoms (N, C, Cα) of the protein chain within 10 Å of the peptide chain are extracted, similarly to the way in RoseTTAFold All-Atom^17^. The kabsch algorithm is then used to align the crystal and predicted protein structures based on these backbone atoms. The same rotation and translation matrices derived from this alignment are applied to superimpose the peptide chains. The pocket-aligned peptide RMSD_all-atom_ quantifies the RMSD of all heavy atoms in the peptide chain, whereas RMSD_unAA_ is calculated exclusively for the heavy atoms of unnatural amino acid residues. The pocket-aligned peptide RMSD_Cα_ follows a similar procedure, and the primary difference is that only the backbone atoms of the protein chains within 10 Å of the Cα atoms in the peptide chain are selected for alignment. Additionally, the RMSD calculation is restricted to the Cα atoms in the peptide chains.

RMSD_all-atom_ is determined by aligning all heavy atoms in the crystal and predicted structures using the kabsch algorithm and subsequently calculating the RMSD across all heavy atoms. When the calculation is limited to unnatural amino acid residues, the metric is denoted as RMSD_unAA_. RMSD_Cα_ is derived by aligning the Cα atoms from the crystal and predicted structures and computing the RMSD between these atoms exclusively.

#### 2.5.2 Structure assessments

The structures were independently evaluated using the online validation server MolProbity^37^, which is a structural analysis tool that provides information on macromolecular accuracy by assessing the quality of the structure based on atomic contact analysis, geometry, and backbone torsion angles. This tool is commonly used for validating the overall structural integrity and stereochemistry of macromolecules.

## 3. Result and Discussion

Building upon the AlphaFold-Multimer framework, we developed a model capable of accurately predicting the three-dimensional structures of peptide monomers and complexes containing unnatural amino acids. During the construction of multiple sequence alignments (MSA), all unnatural amino acids were initially treated as the unknown residue (X). To overcome this limitation, we integrated atom-level embedding information specific to peptides into the AlphaFold-Multimer model, as shown in Fig. 1B. This enhancement allows the model to distinguish among various unnatural amino acid residues while enabling multi-scale modeling of peptide molecules. Since AlphaFold-Multimer models each atom by predicting seven torsion angles for every amino acid along with their predefined rigid groups and atomic coordinates, we extended this methodology to unnatural amino acids. Specifically, we defined corresponding rigid groups and initial atomic coordinates for these residues, as shown in Fig. 1C, enabling the model to predict their atomic coordinates in an end-to-end manner. Next, we fine-tuned the original five AlphaFold-Multimer parameter sets using linear peptides containing unnatural amino acids, as illustrated in Fig. 1A. Furthermore, by modifying the relative position encoding matrix, we expanded this model’s capabilities to predict the structures of cyclic peptides with unnatural residues, as shown in Fig. 1D. To enhance the model further, we parameterized the unnatural amino acids, allowing for the relaxation of spatial structures containing these unnatural residues, as illustrated in Fig. 1E. To the best of our knowledge, this work represents the first effort to fine-tune AlphaFold-Multimer specifically for the structure prediction of cyclic peptides with unnatural amino acids, more details about the workflow can be seen in Methods and Materials section.

**Figure 1.**
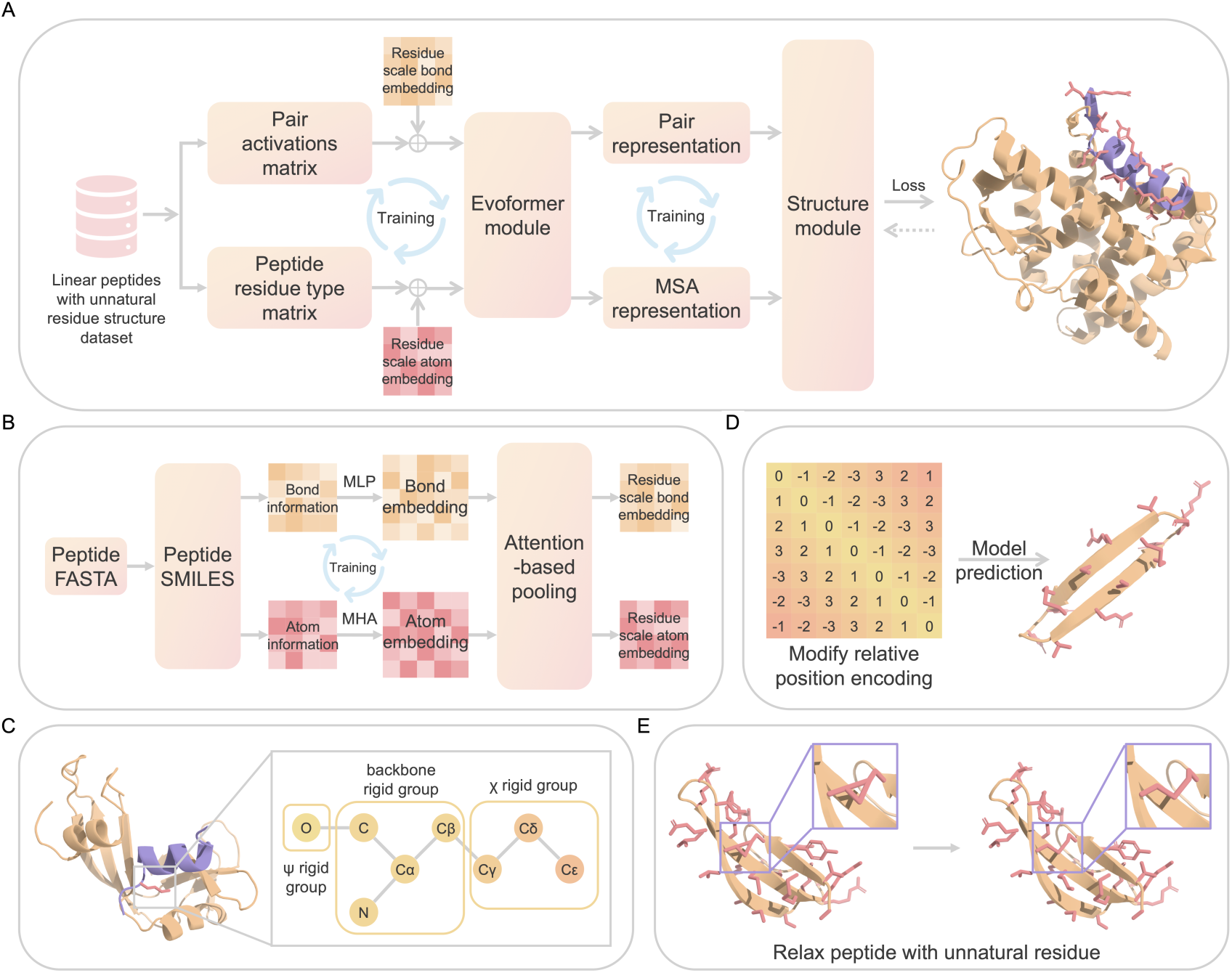
Overview of HighFold2. (A) The training process of structure prediction model. (B) The architecture of the atomic-scale feature extraction module for peptides. MLP and MHA represent the multi-layer perceptron and multi-head attention, respectively. (C) The definition of rigid groups for unnatural amino acids. (D) The method for modifying the relative position encoding in the prediction model to enable the prediction of cyclic peptide monomers and complexes containing unnatural amino acids. (E) The relaxation of three-dimensional structures containing unnatural amino acids to eliminate spatial clashes.

### 3.1 Accurately predict the structure of cyclic peptides with unnatural amino acids

HighFold2 can accurately predict the three-dimensional structures of cyclic peptide monomers containing unnatural amino acids and their complexes with proteins. Since we trained five distinct models, five predicted structures can be generated during testing. The model automatically ranks these predicted structures according to predefined criteria, and we only select the rank one structure for comparison with the native structure. As shown in Fig. 2A, the model achieves a median RMSD_Cα_ (for complexes, RMSD_Cα_ refers to pocket-aligned peptide RMSD_Cα_) of 1.891 Å on the test set of cyclic peptides with unnatural amino acids. Most samples have RMSD_Cα_ values near 2 Å, with 20 out of 34 test samples having RMSD_Cα_ less than 2 Å. This suggests the model’s predictions of cyclic peptide backbones containing unnatural amino acids are highly accurate. The median RMSD_all-atom_ (for complexes, RMSD_all-atom_ refers to pocket-aligned peptide RMSD_all-atom_) is 2.872 Å, with most samples clustering around 3 Å. Six samples had RMSD_all-atom_ values under 2 Å, indicating the model’s reasonable ability to predict side-chain conformations. Given that side-chain positions are not fixed in real scenarios but can undergo torsional adjustments, the model’s performance for RMSD_all-atom_ is considered acceptable. To evaluate the model’s precision in predicting structures of unnatural amino acids, we computed RMSD_unAA_ (for complexes, RMSD_unAA_ refers to pocket-aligned peptide RMSD_unAA_), which yielded a median value of 2.579 Å. This indicates that the model accurately predicts most unnatural amino acid residues. Fig. 2B shows all RMSD distributions in this test set, and the detailed predictions for each unnatural cyclic peptide sample can be found in Table S2.

Like AlphaFold-Multimer, HighFold2 provides a prediction confidence score for each amino acid residue. To investigate the relationship between these confidence scores and RMSD_Cα_, RMSD_all-atom_, and RMSD_unAA_, we obtained the predicted local-distance difference test for peptides (peptide pLDDT) and the predicted local-distance difference test for unnatural amino acid residues (unAA pLDDT). For complexes, peptide pLDDT is the average pLDDT score of the peptide ligand, while for peptide monomers, it is the average pLDDT score of the entire structure. As shown in Fig. 2C and Fig. 2D, peptide pLDDT is linearly correlated with both RMSD_Cα_ and RMSD_all-atom_. As peptide pLDDT increases, the RMSD values between predicted and true structures decrease. In our test set of cyclic peptides with unnatural amino acids, the coefficient of determination (R²) between peptide pLDDT and RMSD_Cα_ is 0.456, while the R² between peptide pLDDT and RMSD_all-atom_ is 0.537. This indicates that peptide pLDDT can serve as a useful metric for roughly assessing the accuracy of predictions for cyclic peptides containing unnatural amino acids, both in monomeric form and as peptide ligands. Fig. 2E demonstrates that unAA pLDDT is also linearly correlated with RMSD_unAA_, with an R² value of 0.377. Although the correlation is the weakest among the three scenarios, this still indicates that unAA pLDDT can serve as a valuable indicator for evaluating the prediction accuracy of unnatural amino acids. The details of peptide pLDDT and unAA pLDDT can be seen in Table S3.

We also visualized the model’s predicted structures for cyclic peptide monomers containing unnatural amino acids and complexes with this kind of cyclic peptide ligands. Fig. 2F shows the predicted structure of mutant human alpha-defensin 1 (PDB ID 3LO6), where the RMSD_Cα_ is as low as 0.643 Å and the RMSD_all-atom_ is 1.254 Å. The predicted position of the unnatural amino acid alpha-aminobutyric acid closely matches that in the crystal structure, achieving an RMSD_unAA_ of 0.426 Å. Notably, this structure is cyclized through three pairs of disulfide bonds, which the model also successfully predicted. Fig. 2G presents the predicted structure of a complex between HIV-1 integrase and a cyclic peptide containing an unnatural amino acid (PDB ID 3WNG). The model not only accurately predicted the binding site between the cyclic peptide ligand and the protein but also predicted the cyclic peptide’s spatial structure. The pocket-aligned peptide RMSD_Cα_ is 1.166 Å, and the pocket-aligned peptide RMSD_all-atom_ is 1.701 Å. This cyclic peptide contains a D-proline residue, and the model’s prediction for this unnatural amino acid is also close to the crystal structure’s position. Furthermore, the model accurately predicted the head-to-tail cyclization feature of this peptide.

**Figure 2.**
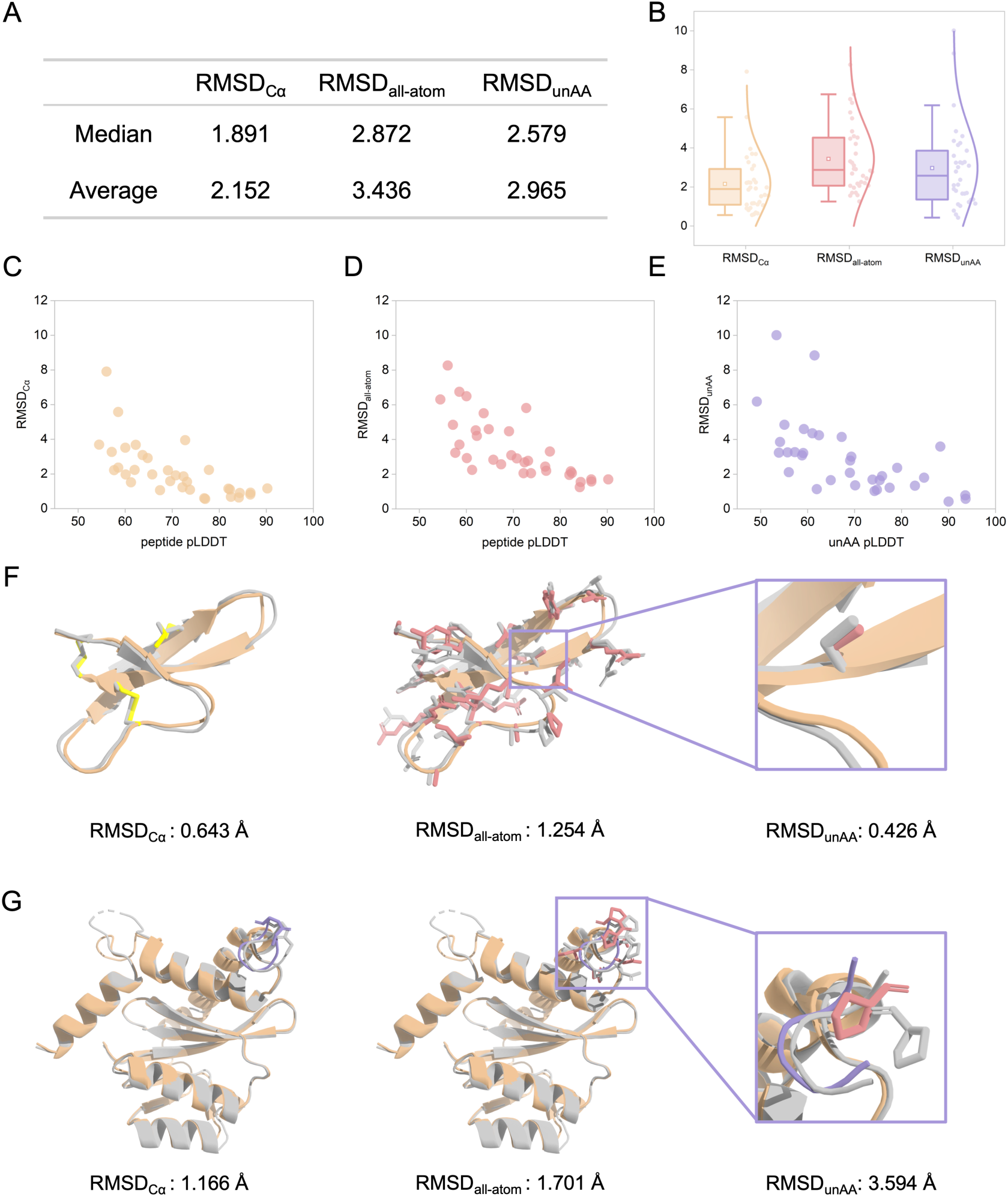
The performance of predicting the structure of cyclic peptide monomers and complexes containing unnatural amino acids. (A) The overall accuracy of the model in the independent cyclic peptide test set. (B) The distributions of all RMSD for cyclic peptide samples. (C) The correlation between peptide pLDDT and RMSD_Cα_ in the independent cyclic peptide test set. (D) The correlation between peptide pLDDT and RMSD_all-atom_ in the independent cyclic peptide test set. (E) The correlation between unAA pLDDT and RMSD_unAA_ in the independent cyclic peptide test set. (F) An example of the prediction for the cyclic peptide monomer containing the unnatural amino acid. (G) An example of the prediction for the cyclic peptide complex containing the unnatural amino acid. The native structure is colored grey, the predicted structure is colored orange, the cyclic peptide in the predicted complex is shown in purple, disulfide bonds are highlighted in yellow, and the side chains are emphasized in red.

### 3.2 Ablation experiments

We conducted a series of ablation studies to elucidate the contributions of individual modules in HighFold2 to the accuracy of predicting cyclic peptide structures with unnatural residues. We systematically removed the ensemble, modifications of the relative position encoding matrix (MRPE), and atomic-scale feature extraction (AFE) from the original model. Then, we evaluated its performance on the test set of cyclic peptides. The results, shown in Fig. 3A, indicate that the full model achieved the lowest average RMSD_Cα_ (2.152 Å), average RMSD_all-atom_ (3.436 Å), and average RMSD_unAA_ (2.965 Å), underscoring the indispensability of these three components. Detailed ablation results for each test sample are provided in Tables S4, S5, and S6.

During testing on cyclic peptides, the model generates five predicted structures for each sample and ranks them using AlphaFold-Multimer’s original criteria. When predictions relied solely on the first parameter set, prediction accuracy diminished, with the average RMSD_Cα_ increasing by 0.266 Å, the average RMSD_all-atom_ by 0.164 Å, and the average RMSD_unAA_ by 0.258 Å, as illustrated in Fig. 3B. This highlights the effectiveness of the ensemble approach. Reliance on a single parameter set tends to significant structural errors for certain samples, such as the β-sheet structure in 2M1P, which was mispredicted, as visualized in Fig. 3C.

Building on the model without the ensemble module, we further removed modifications to the relative position encoding matrix. By modifying the relative position encoding matrix, we ensured the precise representation of the relative positions of cyclized residues, allowing accurate modeling of their connectivity. Without this adjustment, prediction performance deteriorated markedly. As shown in Fig. 3D, many cyclic peptide samples were erroneously predicted as linear peptides, leading to significant increases in all RMSD. The largest increase for RMSD_unAA_, 8.594 Å, occurred for 1T9E, as shown in Fig. 3E, where the model failed to predict its head-to-tail cyclization.

Then, we evaluated the removal of the atomic-scale feature extraction module, which integrates atomic-level peptide features with AlphaFold-Multimer’s residue-level representation to enable multi-scale peptide modeling. Compared to predictions made using only the first fine-tuned parameter set, all RMSD metrics showed a slight increase, as illustrated in Fig. 3F, indicating that the atomic-scale feature extraction module contributes to improving the structural predictions of cyclic peptides. Fig. 3G visualizes the comparison between the predicted structure from this ablation model and the native structure (PDB ID 2MSQ). The overall differences are substantial, with significant deviations in the predicted positions of the disulfide bonds.

**Figure 3.**
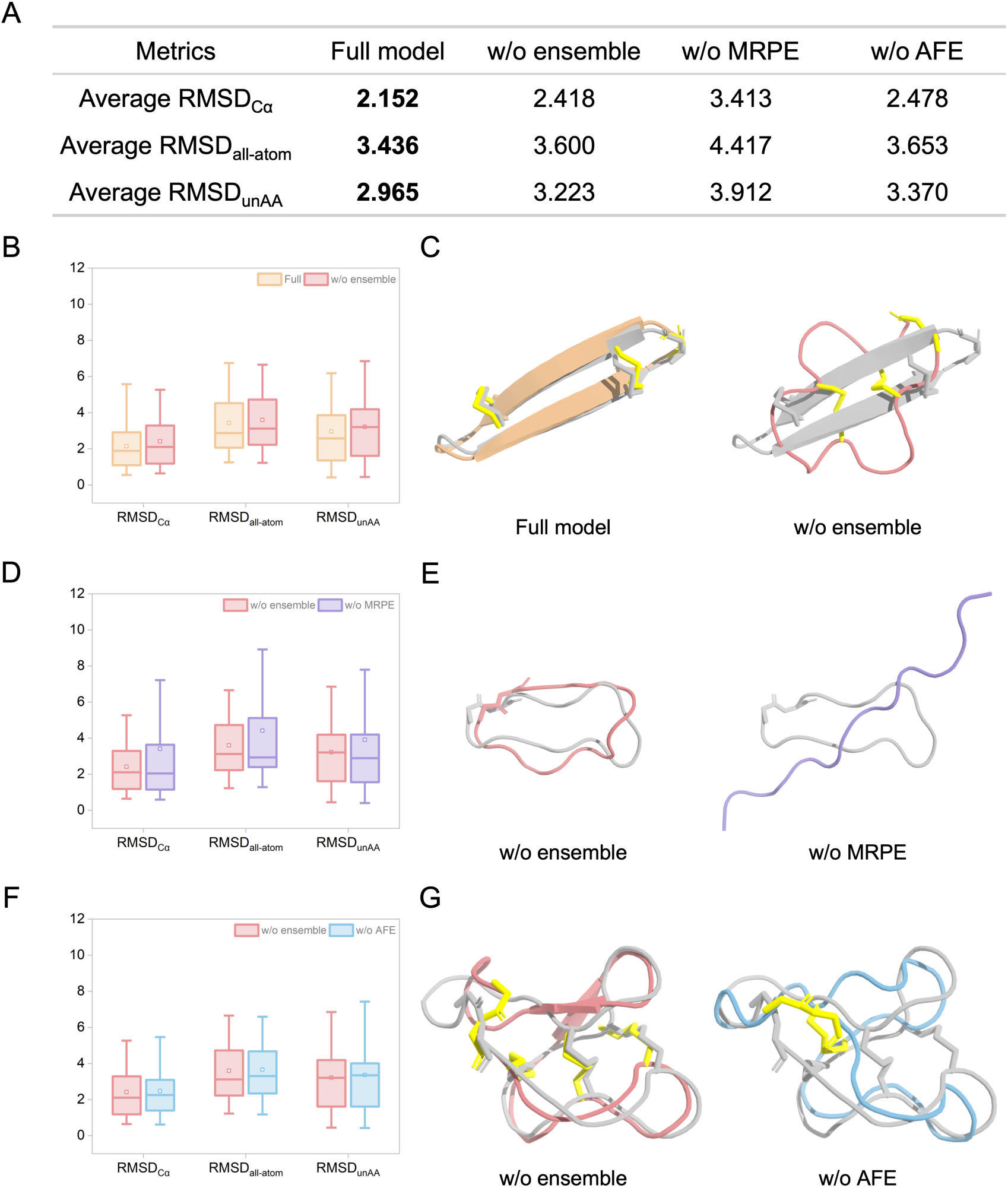
The analysis of the ablation experiments. (A) Predictive performance comparison between the full model and its variants on the independent cyclic peptide test set. w/o stands for without. MRPE and AFE represent the modified relative position encoding and atomic-scale feature extraction, respectively. (B) The comparison of distributions between the full model and the model without the ensemble. (C) A comparison of the prediction for the cyclic peptide containing the unnatural amino acid between the full model and the model without the ensemble. (D) The comparison of distributions between the model without the ensemble and the model without the modified relative position encoding. (E) A comparison of the prediction for the cyclic peptide containing the unnatural amino acid between the model without the ensemble and the model without the modified relative position encoding. (F) The comparison of distributions between the model without the ensemble and the model without the atomic-scale feature extraction. (G) A comparison of the prediction for the cyclic peptide containing the unnatural amino acid between the model without the ensemble and the model without the atomic-scale feature extraction.

### 3.3 Accurately predict the structure of linear peptides with unnatural amino acids

We also evaluated the model’s predicted performance on the test set of linear peptides containing unnatural amino acids. As shown in Fig. 4A, HighFold achieved a median RMSD_Cα_ of 0.994 Å, a median RMSD_all-atom_ of 1.906 Å, and a median RMSD_unAA_ of 1.971 Å on the linear peptide test set. These results confirm the model’s strong ability to predict the three-dimensional structures of linear peptide monomers and their complexes involving unnatural amino acids. Detailed prediction results for each test sample are available in Table S7. Compared to its performance on cyclic peptides, the model performed better on linear peptides. This improved performance likely arises from the training data being based on linear peptide structures. We hypothesize that as more cyclic peptide structures are resolved in the PDB database, incorporating such data into the training process will further enhance the model’s predictive accuracy for cyclic peptides with unnatural amino acids.

In the linear peptide test set, the peptide pLDDT also exhibited a linear relationship with both RMSD_Cα_ and RMSD_all-atom_. As shown in Fig. 4C and Fig. 4D, the R² between peptide pLDDT and RMSD_Cα_ is 0.372, while the R² between peptide pLDDT and RMSD_all-atom_ is 0.397. Notably, the linear relationship between peptide pLDDT and RMSD_all-atom_ is stronger than that with RMSD_Cα_, consistent with observations in the cyclic peptide test set. This suggests that peptide pLDDT reflects not only the accuracy of the predicted backbone but also the side chain accuracy. The R² between unAA pLDDT and RMSD_unAA_ is 0.351, also showing the weakest correlation like in the cyclic peptide dataset. Table S8 shows the detailed peptide pLDDT and unAA pLDDT.

Fig. 4F and Fig. 4G visualize the predicted structures of a linear peptide monomer and a peptide-protein complex, respectively. Fig. 4F shows the predicted structure of the NK1 agonist Phyllomedusin, which includes a pyroglutamic acid residue (PDB ID 2NOR). The predicted structure achieved the RMSD_Cα_ of 0.480 Å, indicating high accuracy in backbone prediction. However, the RMSD_all-atom_ is 1.845 Å, reflecting some inaccuracies in the side chain predictions. Fig. 4G illustrates the predicted structure of a peptide-protein complex containing an alpha-aminoisobutyric acid residue (PDB ID 1FEV). The predicted structure had the RMSD_Cα_ of 0.387 Å and the RMSD_unAA_ of 0.474 Å, indicating not only accurate backbone prediction but also accurate predictions for the unnatural residue.

**Figure 4.**
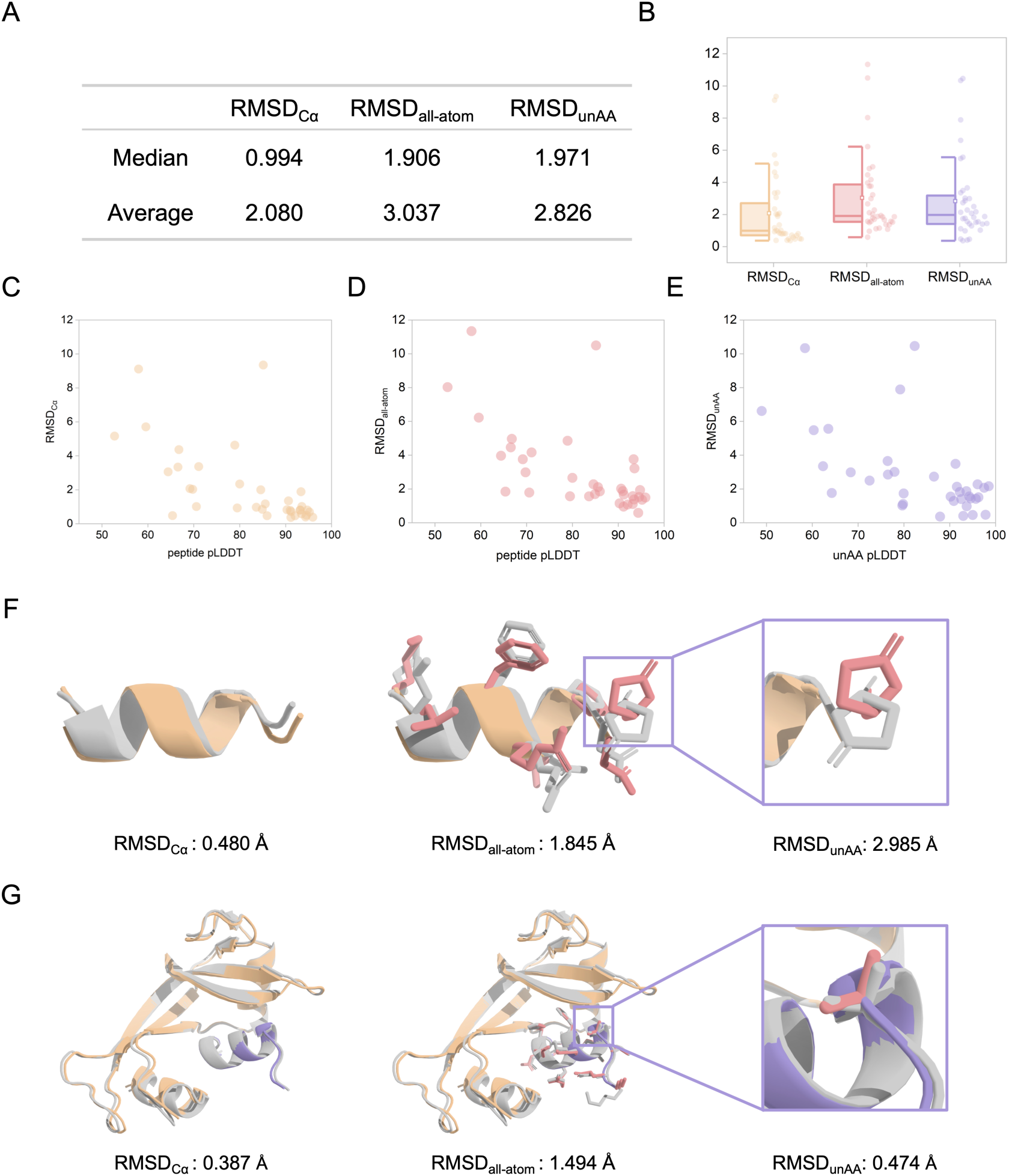
The performance of predicting the structure of linear peptide monomers and complexes containing unnatural amino acids. (A) The overall accuracy of the model in the linear peptide test set. (B) The distributions of all RMSD for linear peptide samples. (C) The correlation between peptide pLDDT and RMSD_Cα_ in the linear peptide test set. (D) The correlation between peptide pLDDT and RMSD_all-atom_ in the linear peptide test set. (E) The correlation between unAA pLDDT and RMSD_unAA_ in the linear peptide test set. (F) An example of the prediction for the linear peptide monomer containing the unnatural amino acid. (G) An example of the prediction for the linear peptide complex containing the unnatural amino acid.

### 3.4 Relaxation of structure with unnatural residues

In this work, based on the efficient relaxation steps from AlphaFold-Multimer, we introduce an additional set of force field parameters for unnatural amino acids, which are obtained through ab initio calculations and compatible with the Amber force field. These parameters aim to improve the accuracy of the local geometry of the predicted models. The improvements include optimizing protein folding accuracy, reducing clashes, and enhancing the stereochemical correctness of protein residues.

Force field parameters for unnatural amino acids are essential for the relaxation process. When custom parameters are not available, the reliability of relaxation is limited. While general force fields like GAFF or CGenFF^38^ can assign parameters to various compounds, their lack of specific optimization for unnatural amino acids often results in inaccurate structural predictions. Fig. 5A shows that after 6-31G* structural optimization, the unnatural amino acids are highly consistent with the crystal structures, with an average RMSD of 0.153 Å, significantly improving the correlation between the unnatural amino acid conformations and the real structures.

Subsequently, we generated topology and coordinate files for the target structure in tleap. By modifying the OpenMM input files, we overcame the limitation of tleap’s inability to output multichain systems, enabling the automated relaxation process for more complex systems. As shown in Fig. 5B, after relaxation, the quality of the structures for all unnatural residues improved significantly compared to the results from the structural prediction models. Table S9 shows that there were minimal or no spatial clashes, unfavorable rotamers, or abnormal Ramachandran values, with most residues in favorable regions in the independent cyclic peptide test set. In Fig. 5C, due to the lack of consideration for reasonable atomic contacts, some non-bonded atomic pairs in the predicted structure were found to be too close. These errors were corrected through an energy minimization procedure without significantly increasing computational costs. Even though the prediction model provided a high-quality overall structure, some side-chain deviations and unreasonable geometry were still observed, suggesting further molecular dynamics simulations or quantum mechanics/molecular mechanics (QM/MM) calculations to improve residue stereochemistry.

To further evaluate the optimization effect, we selected the covalent bonds between the natural amino acid portion and the modified group in unnatural amino acids as an evaluation criterion. Since bond lengths in the predicted models are based on constant, they cannot be adjusted according to environmental changes. Table S10 shows that after relaxation, the bond lengths of the unnatural amino acids are closer to those in the crystal structures^39^. Finally, we analyzed the impact of the relaxation process on the overall RMSD values. As shown in Table S11 and Table S12, the RMSD_Cα_, RMSD_all-atom_, and RMSD_unAA_ values remain little changed for the cyclic and linear peptide test sets. Although side-chain adjustments introduced some local changes, they did not negatively affect the overall accuracy of the structural predictions.

We corrected unreasonable regions in the predicted structures through relaxation, significantly improving their geometric quality and accuracy. This process reduced the need for manual intervention and enhanced the efficiency and precision of handling systems containing unnatural amino acids.

**Figure 5.**
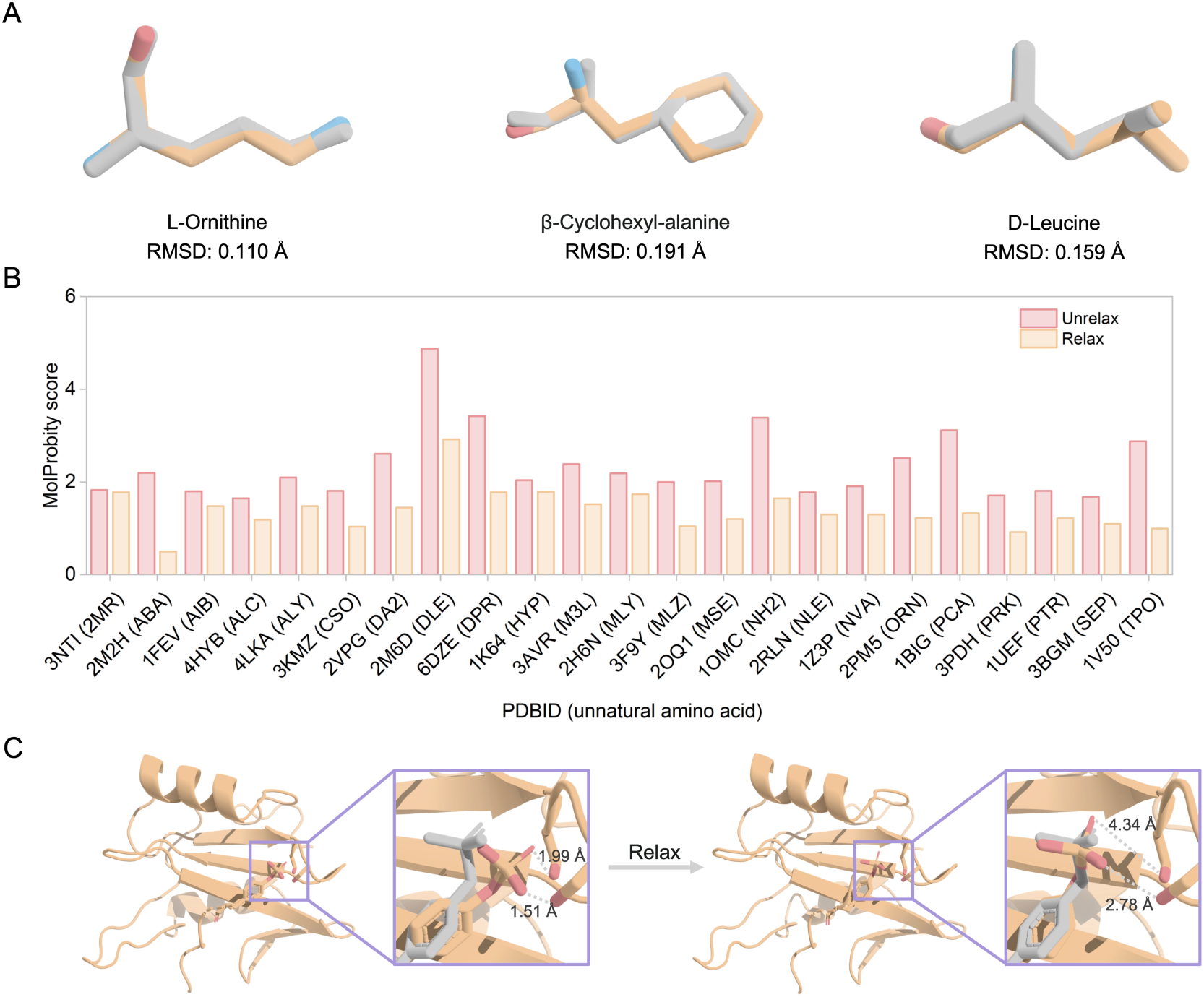
The relaxation for the predicted structure. (A) Structural alignment between the unnatural amino acid after QM (B3LYP/6-31G*) optimization and crystal structure. (B) Quality improvement in structures containing unnatural amino acids after relaxation. MolProbity integrates multiple evaluation criteria, with lower scores indicating better structure quality. (C) The comparison between the structures before and after relaxation for PDB ID 2CI9 containing O-phosphotyrosine. The crystal structure is shown in grey, the predicted structure is shown in orange, the N atom is shown in blue, and the O atom is shown in red.

## 4. Conclusion

In this work, we developed a method called HighFold2, based on AlphaFold-Multimer, capable of accurately predicting the three-dimensional structures of cyclic peptide monomers containing unnatural amino acids and their complexes. To adapt AlphaFold-Multimer for the prediction of structures involving unnatural amino acids, we added a neural network module to characterize the atom-level properties of peptides. This not only enabled the distinction between different unnatural amino acids but also facilitated the multi-scale modeling of peptide molecules. We also extended the predefined rigid groups and initial atomic coordinate information for natural amino acids in AlphaFold-Multimer to encompass unnatural amino acids, successfully enabling the prediction of their structures. Given the scarcity of cyclic peptide structural data, we employed a zero-shot learning strategy, initially training the model using linear peptide data containing unnatural amino acids. Then, by modifying the relative position encoding matrix in the model, we achieved accurate predictions of cyclic peptides containing unnatural amino acids. Furthermore, by combining Gaussian, AmberTools, and OpenMM, we developed an efficient relaxation workflow, significantly reducing spatial clashes and improving geometric accuracy. Further testing demonstrated that HighFold2 showed outstanding prediction performance on both independent cyclic peptide and linear peptide test sets, and various ablation experiments validated the effectiveness of its modifications.

## Data availability

All data this work uses is available at https://github.com/hongliangduan/HighFold2 or the Supporting Information.

## Code availability

The code of this work is available at https://github.com/hongliangduan/HighFold2.

## Acknowledgment

This work was supported by the Natural Science Foundation of Zhejiang Province (LD22H300004).

## Author contributions

Cheng Zhu: conceptualization; data curation; methodology; software; writing - original draft. Sen Cao: methodology; writing - original draft. Tianfeng Shang: supervision. Hongliang Duan: supervision. All authors: writing - review and editing.

