## Supporting Information for "Predicting the structures of cyclic peptides containing unnatural amino acids by HighFold2"

**Section S1. Datasets**

**Linear peptides with unnatural amino acids dataset**

The PDB ID of the linear peptide training set follows: 3iqj, 3bh9, 3da9, 4i7b, 4bl0, 3bhb, 3bzi, 3k27, 4igk, 2gfa, 3pxe, 3dpq, 4hpt, 2h2h, 1q1a, 3tu7, 3cu8, 1s5p, 3fqr, 1qja, 1ywt, 4apj, 3wdz, 4j6s, 4bg6, 2od2, 3coj, 1qjb, 4fl5, 3n9p, 1umw, 3os5, 1p22, 4hy9, 3uat, 3axy, 2rfi, 1gxc, 3ubw, 3lw1, 3f9x, 4ezh, 3uvw, 2lax, 3al3, 3n9q, 2p5b, 4k45, 3hna, 3d9p, 3pfv, 4buz, 1i8h, 3d9m, 3glr, 3bux, 3p36, 1jlu, 3fxx, 3ual, 3bum, 2h6k, 1j4l, 1o9s, 2iui, 3lqi, 2bbu, 3a0a, 4hkc, 2lkj, 4bu1, 3omh, 3bh8, 2vpe, 2lb0, 3qo2, 2g6q, 4mzf, 4au7, 1lqb, 3bu6, 3tzd, 2rmx, 4gvc, 4jxt, 1lm8, 2w5z, 3q1i, 2v7d, 2x4y, 2ror, 1y98, 3buw, 2os2, 2lue, 2lnw, 4iea, 1t2v, 1t29, 2x4w, 3v7d, 3ob1, 1py1, 2lo6, 1fgl, 2ot7, 2pld, 4ch2, 2cef, 1p13, 2ox0, 3wp0, 4hon, 2ovr, 2jkt, 1f1w, 2lb3, 4o42, 1nex, 4n4i, 2jkr, 2oq6, 2c1n, 3cbm, 2l5j, 3u4s, 2jmj, 1i8g, 3u5p, 3nf3, 3d9k, 3iiy, 2c1j, 4kv1, 2q8c, 2a0t, 2ovq, 1nzs, 1v4f, 2jnd, 3alo, 4gl9, 3d9l, 2mc1, 2ast, 2z8p, 1qg1, 2jqc, 2cez, 2ybp, 3d4b, 2h4f, 2hwl, 1yc5, 2lyw, 2mjv, 2k17, 1bmb, 2m7j, 4gfu, 1shc, 2rt5, 2cfj, 3qzs, 2yu7, 3ueo, 3u5o, 3buo, 2l3r, 1tce, 2ght, 2lct, 2l11, 1g1h, 2wp2, 2l1b, 2ghq, 4c78, 2lsp, 3swc, 1ju5, 1oo4, 2q8e, 2fuu, 2lgk, 1f3r, 2nmb, 2lvm, 2qqs, 3uvy, 1fpr, 2kvm, 2e3k, 4b9w, 3dpc, 2kwn, 3mxc, 4dow, 1guw, 3uzd, 2m3m, 3n9n, 3mxy, 1eeo, 2pie, 4edu, 2q8y, 1yrk, 4l1u, 1jm4, 1fhr, 3olr, 3omc, 2rs9, 3ask, 3wp1, 1fu5, 2g46, 1sps, 4jfz, 4hcz, 1yvh, 3vrp, 1cfn, 2rny, 3zni, 2rsn, 1ma3, 1kc2, 2w3o, 1mw4, 4gag, 1f58, 1lck, 1fbv, 3ob2, 4jg1, 1g6g, 2vif, 2jql, 4nw2, 1j4q, 4ihl, 2jqi, 2rnw, 2v87, 2m0o, 3dpo, 1jsp, 2lto, 4glr, 3maz, 1j4p, 2rnx, 3sqd, 3o34, 1aot, 2ktb, 3va4, 2v89, 1a81, 4fj3, 3a08, 2dvr,

2d3h, 2wh0, 2pqw, 3a0m, 3abn, 3ai6, 2rje, 3a1h, 2dvq, 1pua, 1v6d, 1dzi, 1f8a, 3kv4, 4jqi, 4bj3, 1p16, 2y5t, 4bu0, 4bkl, 4jg0, 4lor, 4ft2.

The validation set of the linear peptides follows: 2x4x, 3i91, 4llb, 1aya, 1ayb, 1pdq, 4iur, 3mhr, 3zkf, 1lf8, 1d4w, 3g7l, 1pfb, 3tl0, 4jiz, 1jyr, 3dm1, 3g2v, 3i90, 3unn, 2hdx, 3g2t, 1ayc, 3poa, 2gfr, 2g35, 3bun, 3d9n, 2bqz, 3uvx, 2rr4, 2pnx, 2iuh, 2wp1, 2mg5, 2laz, 2lb2, 2mfq, 1t15, 1irs, 3r93, 2laj, 3svm, 2lay, 3dpp, 3ml4, 1z3l, 1z3m, 4ajy, 2npv, 1jy6, 1tze, 3muk, 2v88, 2v86, 1ai1, 1n0x.

The test set of the linear peptides follows: 2zol, 4hyb, 3l6f, 2zok, 3kmt, 2zsw, 2rln, 3fqx, 3avr, 1rbd, 3fqu, 4lka, 3u5n, 1z3p, 2ci9, 3bgm, 1fev, 1uef, 3f9y, 2oq1, 1mxq, 2nor, 1jy9, 2h6n, 3kmz, 4jaa, 3cmh, 3ejh, 1v50, 2vpg, 2k7l, 6cmh, 3pdh, 3nti, 2fx8, 3idj.

**Cyclic peptides with unnatural amino acids dataset**

The independent cyclic peptide test set follows: 2ew4, 2msq, 6dz9, 1t9e, 2m2s, 2pm5, 4e86, 1big, 3wng, 2m2x, 5ug3, 4e83, 2m6d, 2crd, 1p9g, 2mg6, 1omc, 2m2g, 2n8e, 2j15, 6dze, 6dza, 2m7j, 2m2h, 5n99, 6dzc, 2m1p, 2m62, 2mfx, 2ih6, 6my2, 1k64, 6dzb, 3lo6.

### Section S2. Rigid group assignments

**Table S1.** The rigid group assignments for unnatural amino acids.

| Residue | Backbone | $\psi$ | $\chi_1$ | $\chi_2$ | $\chi_3$ | $\chi_4$ |
|---|---|---|---|---|---|---|
| 2MR | N, $C_\alpha$, C, $C_\beta$ | O | $C_\gamma$ | $C_\delta$ | $N_\varepsilon$ | $C_\zeta$, $N_{\eta1}$, $N_{\eta2}$, $C_{Q1}$, $C_{Q2}$ |
| ABA | N, $C_\alpha$, C, $C_\beta$ | O | $C_\gamma$ | - | - | - |
| AIB | N, $C_\alpha$, C, $C_{\beta1}$, $C_{\beta2}$ | O | - | - | - | - |
| ALC | N, $C_\alpha$, C, $C_\beta$ | O | $C_\gamma$ | $C_{\delta1}$, $C_{\delta2}$, $C_{\varepsilon1}$, $C_{\varepsilon2}$, $C_\zeta$ | - | - |
| ALY | N, $C_\alpha$, C, $C_\beta$ | O | $C_\gamma$ | $C_\delta$ | $C_\varepsilon$ | $N_\zeta$, $O_\eta$, $C_\eta$, $C_{\eta3}$ |
| CSO | N, $C_\alpha$, C, $C_\beta$ | O | $S_\gamma$ | $O_\delta$ | - | - |
| DA2 | N, $C_\alpha$, C, $C_\beta$ | O | $C_\gamma$ | $C_\delta$ | $N_\varepsilon$ | $C_\zeta$, $N_{\eta1}$, $N_{\eta2}$, $C_1$, $C_2$ |
| DLE | N, $C_\alpha$, C, $C_\beta$ | O | $C_\gamma$ | $C_{\delta1}$, $C_{\delta2}$ | - | - |
| DPR | N, $C_\alpha$, C, $C_\beta$, $C_\gamma$, $C_\delta$ | O | - | - | - | - |

| | | | | | | |
|---|---|---|---|---|---|---|
| HYP | N, $C_\alpha$, C, $C_\beta$, $C_\gamma$, $C_\delta$ | O | $O_{\delta 1}$ | - | - | - |
| M3L | N, $C_\alpha$, C, $C_\beta$ | O | $C_\gamma$ | $C_\delta$ | $C_\varepsilon$ | $N_\zeta$, $C_{M1}$, $C_{M2}$, $C_{M3}$ |
| MLY | N, $C_\alpha$, C, $C_\beta$ | O | $C_\gamma$ | $C_\delta$ | $C_\varepsilon$ | $N_\zeta$, $C_{\eta 1}$, $C_{\eta 2}$ |
| MLZ | N, $C_\alpha$, C, $C_\beta$ | O | $C_\gamma$ | $C_\delta$ | $C_\varepsilon$ | $N_\zeta$, $C_M$ |
| MSE | N, $C_\alpha$, C, $C_\beta$ | O | $C_\gamma$ | Se | $C_\varepsilon$ | - |
| NH2 | N | - | - | - | - | - |
| NLE | N, $C_\alpha$, C, $C_\beta$ | O | $C_\gamma$ | $C_\delta$ | $C_\varepsilon$ | - |
| NVA | N, $C_\alpha$, C, $C_\beta$ | O | $C_\gamma$ | $C_\delta$ | - | - |
| ORN | N, $C_\alpha$, C, $C_\beta$ | O | $C_\gamma$ | $C_\delta$ | $N_\varepsilon$ | - |
| PCA | N, $C_\alpha$, C, $C_\beta$, $C_\gamma$, $C_\delta$, $O_\varepsilon$ | O | - | - | - | - |
| PRK | N, $C_\alpha$, C, $C_\beta$ | O | $C_\gamma$ | $C_\delta$ | $C_\varepsilon$ | $N_\zeta$, $C_{AL}$, $O_{AD}$, $C_{AF}$, $C_{AA}$ |
| PTR | N, $C_\alpha$, C, $C_\beta$ | O | $C_\gamma$ | $C_{\delta 1}$, $C_{\delta 2}$, $C_{\varepsilon 1}$, $C_{\varepsilon 2}$, $C_\zeta$, $O_\eta$ | P | $O_{1P}$, $O_{2P}$, $O_{3P}$ |

| | | | | | | |
|---|---|---|---|---|---|---|
| SEP | N, $C_\alpha$, C, $C_\beta$ | O | $O_\gamma$ | P | $O_{1P}$, $O_{2P}$, $O_{3P}$ | - |
| TPO | N, $C_\alpha$, C, $C_\beta$ | O | $O_{\gamma1}$, $C_{\gamma2}$ | P | $O_{1P}$, $O_{2P}$, $O_{3P}$ | - |

### Section S3. Prediction performance for cyclic peptide test set

**Table S2.** All RMSD for the independent cyclic peptide test set.

| PDB ID | $RMSD_{C\alpha}$ | $RMSD_{all\text{-}atom}$ | $RMSD_{unAA}$ |
|---|---|---|---|
| 2ew4 | 1.092 | 2.058 | 1.889 |
| 2msq | 1.861 | 2.684 | 2.081 |
| 6dz9 | 2.211 | 4.472 | 2.790 |
| 1t9e | 1.971 | 2.926 | 2.115 |
| 2m2s | 2.234 | 3.307 | 2.368 |
| 2pm5 | 0.902 | 1.553 | 1.801 |
| 4e86 | 0.814 | 1.700 | 0.783 |
| 1big | 1.153 | 1.959 | 1.109 |
| 3wng | 1.166 | 1.701 | 3.594 |
| 2m2x | 3.517 | 6.496 | 4.358 |
| 5ug3 | 3.270 | 4.846 | 3.242 |
| 4e83 | 0.920 | 1.579 | 0.585 |
| 2m6d | 1.921 | 2.910 | 3.009 |
| 2crd | 1.104 | 2.165 | 1.032 |
| 1p9g | 1.230 | 2.060 | 4.851 |
| 2mg6 | 1.519 | 2.250 | 3.257 |
| 1omc | 1.062 | 2.571 | 1.657 |
| 2m2g | 0.560 | 2.197 | 1.658 |
| 2n8e | 2.221 | 3.234 | 6.184 |

| | | | |
|---|---|---|---|
| 2j15 | 1.586 | 3.095 | 1.689 |
| 6dze | 2.918 | 4.596 | 3.219 |
| 6dza | 3.950 | 5.816 | 4.147 |
| 2m7j | 3.693 | 6.306 | 3.270 |
| 2m2h | 0.597 | 2.440 | 1.224 |
| 5n99 | 7.907 | 8.265 | 8.844 |
| 6dzc | 3.092 | 5.514 | 4.246 |
| 2m1p | 0.681 | 2.067 | 1.339 |
| 2m62 | 1.975 | 2.834 | 1.358 |
| 2mfx | 3.682 | 4.207 | 4.602 |
| 2ih6 | 2.368 | 3.705 | 3.858 |
| 6my2 | 1.550 | 2.773 | 3.086 |
| 1k64 | 5.576 | 6.748 | 10.010 |
| 6dzb | 2.234 | 4.527 | 1.143 |
| 3lo6 | 0.643 | 1.254 | 0.426 |

**Table S3.** The peptide pLDDT and unAA pLDDT in the independent cyclic peptide test set.

| PDB ID | peptide pLDDT | unAA pLDDT |
|---|---|---|
| 2ew4 | 73.76 | 75.85 |
| 2msq | 72.39 | 69.00 |
| 6dz9 | 69.08 | 69.06 |
| 1t9e | 60.10 | 55.99 |
| 2m2s | 77.79 | 79.07 |
| 2pm5 | 84.34 | 84.79 |
| 4e86 | 86.59 | 93.56 |
| 1big | 81.94 | 74.75 |
| 3wng | 90.22 | 88.25 |
| 2m2x | 60.03 | 61.01 |
| 5ug3 | 57.14 | 53.88 |
| 4e83 | 86.63 | 93.62 |
| 2m6d | 70.74 | 69.31 |
| 2crd | 82.24 | 74.25 |
| 1p9g | 72.18 | 55.06 |
| 2mg6 | 61.23 | 55.73 |
| 1omc | 67.41 | 64.92 |
| 2m2g | 77.00 | 75.34 |
| 2n8e | 57.65 | 49.19 |

| | | |
|---|---|---|
| 2j15 | 69.57 | 73.81 |
| 6dze | 64.77 | 59.06 |
| 6dza | 72.75 | 67.25 |
| 2m7j | 54.46 | 57.28 |
| 2m2h | 76.80 | 77.46 |
| 5n99 | 56.02 | 61.47 |
| 6dzc | 63.70 | 62.41 |
| 2m1p | 82.46 | 82.85 |
| 2m62 | 65.75 | 70.12 |
| 2mfx | 62.24 | 59.20 |
| 2ih6 | 58.52 | 54.12 |
| 6my2 | 73.14 | 58.81 |
| 1k64 | 58.51 | 53.38 |
| 6dzb | 62.01 | 61.94 |
| 3lo6 | 84.15 | 90.00 |

**Section S4. The result of ablation experiments**

**Table S4.** All RMSD for the model without the ensemble in the independent cyclic peptide test set.

| PDB ID | $RMSD_{C\alpha}$ | $RMSD_{all\text{-}atom}$ | $RMSD_{unAA}$ |
|---|---|---|---|
| 2ew4 | 3.301 | 4.392 | 3.855 |
| 2msq | 1.809 | 2.817 | 1.447 |
| 6dz9 | 3.101 | 5.026 | 4.892 |
| 1t9e | 1.991 | 2.967 | 2.190 |
| 2m2s | 1.188 | 2.355 | 1.664 |
| 2pm5 | 0.861 | 1.228 | 1.295 |
| 4e86 | 0.848 | 1.566 | 0.790 |
| 1big | 0.983 | 1.808 | 1.110 |
| 3wng | 1.098 | 1.667 | 3.488 |
| 2m2x | 2.791 | 4.730 | 3.274 |
| 5ug3 | 3.290 | 4.831 | 3.743 |
| 4e83 | 0.958 | 1.662 | 0.532 |
| 2m6d | 2.617 | 3.008 | 4.577 |
| 2crd | 1.110 | 2.128 | 0.649 |
| 1p9g | 1.686 | 2.228 | 5.044 |
| 2mg6 | 1.519 | 2.250 | 3.257 |
| 1omc | 2.366 | 3.450 | 2.964 |
| 2m2g | 1.469 | 2.333 | 1.617 |

| | | | |
|---|---|---|---|
| 2n8e | 2.221 | 3.234 | 6.184 |
| 2j15 | 3.494 | 4.630 | 3.813 |
| 6dze | 2.510 | 4.097 | 3.152 |
| 6dza | 3.445 | 4.941 | 4.195 |
| 2m7j | 3.816 | 6.655 | 2.661 |
| 2m2h | 1.987 | 2.365 | 2.169 |
| 5n99 | 8.268 | 9.957 | 6.854 |
| 6dzc | 3.430 | 6.128 | 3.772 |
| 2m1p | 4.934 | 6.249 | 5.855 |
| 2m62 | 0.949 | 2.206 | 1.864 |
| 2mfx | 2.250 | 2.575 | 4.230 |
| 2ih6 | 2.368 | 3.705 | 3.858 |
| 6my2 | 1.748 | 3.298 | 3.013 |
| 1k64 | 5.267 | 6.369 | 10.272 |
| 6dzb | 1.892 | 4.257 | 0.875 |
| 3lo6 | 0.640 | 1.270 | 0.444 |

**Table S5.** All RMSD for the model without the modified relative position encoding in the independent cyclic peptide test set.

| PDB ID | $RMSD_{C\alpha}$ | $RMSD_{all\text{-}atom}$ | $RMSD_{unAA}$ |
|---|---|---|---|
| 2ew4 | 1.037 | 1.901 | 1.851 |
| 2msq | 2.709 | 3.618 | 2.082 |
| 6dz9 | 4.538 | 6.168 | 2.597 |
| 1t9e | 10.970 | 11.325 | 10.784 |
| 2m2s | 1.283 | 2.894 | 1.555 |
| 2pm5 | 0.912 | 1.620 | 1.353 |
| 4e86 | 0.859 | 1.631 | 0.827 |
| 1big | 1.146 | 2.011 | 1.117 |
| 3wng | 2.137 | 2.915 | 4.201 |
| 2m2x | 12.884 | 13.418 | 11.246 |
| 5ug3 | 3.172 | 4.758 | 4.020 |
| 4e83 | 0.900 | 1.566 | 0.616 |
| 2m6d | 2.324 | 2.938 | 3.604 |
| 2crd | 1.088 | 2.117 | 0.623 |
| 1p9g | 1.672 | 2.520 | 4.913 |
| 2mg6 | 1.542 | 2.395 | 3.447 |
| 1omc | 1.850 | 3.099 | 2.113 |
| 2m2g | 1.640 | 3.282 | 1.558 |
| 2n8e | 7.218 | 8.568 | 13.710 |

| | | | |
|---|---|---|---|
| 2j15 | 1.728 | 2.670 | 1.888 |
| 6dze | 3.576 | 5.116 | 3.761 |
| 6dza | 3.636 | 4.969 | 3.583 |
| 2m7j | 7.620 | 8.389 | 6.258 |
| 2m2h | 1.955 | 2.923 | 1.990 |
| 5n99 | 7.743 | 8.920 | 7.796 |
| 6dzc | 12.494 | 11.963 | 9.198 |
| 2m1p | 0.602 | 2.505 | 0.774 |
| 2m62 | 0.915 | 2.213 | 1.894 |
| 2mfx | 2.273 | 2.425 | 3.877 |
| 2ih6 | 2.564 | 3.846 | 2.361 |
| 6my2 | 1.574 | 2.911 | 3.179 |
| 1k64 | 5.305 | 6.362 | 10.327 |
| 6dzb | 3.592 | 4.951 | 3.499 |
| 3lo6 | 0.596 | 1.284 | 0.404 |

**Table S6.** All RMSD for the model without the atomic-scale feature extraction in the independent cyclic peptide test set.

| PDB ID | $RMSD_{C\alpha}$ | $RMSD_{all\text{-}atom}$ | $RMSD_{unAA}$ |
|---|---|---|---|
| 2ew4 | 3.416 | 4.655 | 4.241 |
| 2msq | 3.121 | 4.443 | 3.235 |
| 6dz9 | 3.025 | 4.677 | 4.006 |
| 1t9e | 1.844 | 2.789 | 2.005 |
| 2m2s | 1.174 | 2.339 | 1.612 |
| 2pm5 | 0.888 | 1.177 | 1.321 |
| 4e86 | 0.902 | 1.675 | 0.776 |
| 1big | 1.079 | 1.909 | 1.194 |
| 3wng | 1.188 | 1.737 | 3.517 |
| 2m2x | 2.573 | 5.087 | 3.177 |
| 5ug3 | 3.481 | 4.926 | 3.777 |
| 4e83 | 0.985 | 1.647 | 0.712 |
| 2m6d | 2.270 | 2.874 | 4.404 |
| 2crd | 1.104 | 2.145 | 0.726 |
| 1p9g | 1.822 | 2.936 | 5.875 |
| 2mg6 | 1.506 | 1.931 | 3.185 |
| 1omc | 2.227 | 3.311 | 2.473 |
| 2m2g | 1.518 | 2.429 | 1.614 |
| 2n8e | 2.335 | 3.334 | 6.237 |

| | | | |
|---|---|---|---|
| 2j15 | 3.362 | 4.463 | 3.622 |
| 6dze | 2.786 | 4.662 | 4.070 |
| 6dza | 3.347 | 4.819 | 4.000 |
| 2m7j | 3.096 | 6.251 | 2.751 |
| 2m2h | 2.240 | 3.295 | 2.589 |
| 5n99 | 8.848 | 8.939 | 8.692 |
| 6dzc | 3.089 | 5.497 | 3.840 |
| 2m1p | 5.469 | 6.594 | 7.434 |
| 2m62 | 1.396 | 2.443 | 1.257 |
| 2mfx | 2.268 | 2.780 | 4.010 |
| 2ih6 | 2.329 | 3.536 | 3.474 |
| 6my2 | 1.666 | 2.921 | 3.455 |
| 1k64 | 5.286 | 6.358 | 9.929 |
| 6dzb | 1.991 | 4.351 | 0.934 |
| 3lo6 | 0.613 | 1.280 | 0.424 |

### Section S5. Prediction performance for linear peptide test set

**Table S7.** All RMSD for the linear peptide test set.

| PDB ID | $RMSD_{C\alpha}$ | $RMSD_{all\text{-}atom}$ | $RMSD_{unAA}$ |
|---|---|---|---|
| 6cmh | 5.708 | 6.222 | 3.349 |
| 2k7l | 3.337 | 4.464 | 5.478 |
| 2fx8 | 0.933 | 1.569 | 1.111 |
| 1fev | 0.387 | 1.494 | 0.474 |
| 1jy9 | 2.342 | 2.659 | 1.737 |
| 2zok | 1.171 | 2.109 | 0.369 |
| 3avr | 0.974 | 1.567 | 2.133 |
| 3pdh | 3.059 | 3.968 | 0.979 |
| 1mxq | 1.015 | 1.789 | 1.764 |
| 2h6n | 5.163 | 8.027 | 10.336 |
| 2ci9 | 0.854 | 1.699 | 1.559 |
| 3kmt | 0.805 | 1.112 | 1.417 |
| 3u5n | 0.789 | 1.875 | 2.731 |
| 2zsw | 1.890 | 3.761 | 1.289 |
| 3nti | 9.113 | 11.341 | 5.559 |
| 1v50 | 2.028 | 2.984 | 2.854 |
| 4lka | 0.826 | 1.152 | 1.823 |
| 3fqu | 0.807 | 1.937 | 3.488 |
| 3idj | 4.631 | 4.853 | 7.891 |

| | | | |
|---|---|---|---|
| 2zol | 0.400 | 0.587 | 0.406 |
| 2oq1 | 1.350 | 2.017 | 2.276 |
| 4jaa | 2.002 | 2.278 | 3.012 |
| 2nor | 0.480 | 1.845 | 2.985 |
| 2rln | 2.071 | 3.762 | 6.615 |
| 4hyb | 0.375 | 0.959 | 1.507 |
| 1uef | 0.716 | 1.667 | 1.549 |
| 1rbd | 0.591 | 1.377 | 1.023 |
| 3fqx | 0.469 | 1.081 | 1.402 |
| 3kmz | 1.017 | 3.221 | 2.070 |
| 3cmh | 3.369 | 4.165 | 2.495 |
| 3ejh | 9.348 | 10.495 | 10.463 |
| 2vpg | 4.367 | 4.966 | 3.655 |
| 3l6f | 0.692 | 1.363 | 2.164 |
| 3bgm | 0.807 | 1.576 | 1.872 |
| 1z3p | 0.503 | 1.524 | 1.440 |
| 3f9y | 0.475 | 1.860 | 0.462 |

**Table S8.** The peptide pLDDT and unAA pLDDT in the linear peptide test set.

| PDB ID | peptide pLDDT | unAA pLDDT |
|---|---|---|
| 6cmh | 59.57 | 62.34 |
| 2k7l | 66.54 | 60.31 |
| 2fx8 | 79.46 | 79.81 |
| 1fev | 95.91 | 97.81 |
| 1jy9 | 79.99 | 79.94 |
| 2zok | 85.57 | 87.88 |
| 3avr | 83.62 | 91.56 |
| 3pdh | 64.41 | 93.69 |
| 1mxq | 70.57 | 64.25 |
| 2h6n | 52.75 | 58.41 |
| 2ci9 | 84.92 | 94.25 |
| 3kmt | 93.16 | 95.69 |
| 3u5n | 90.99 | 86.56 |
| 2zsw | 93.35 | 90.88 |
| 3nti | 57.97 | 63.50 |
| 1v50 | 69.76 | 76.50 |
| 4lka | 90.29 | 92.19 |
| 3fqu | 94.74 | 91.25 |
| 3idj | 78.93 | 79.19 |
| 2zol | 94.35 | 92.94 |

| | | |
|---|---|---|
| 2oq1 | 90.62 | 96.07 |
| 4jaa | 84.59 | 77.94 |
| 2nor | 65.38 | 68.38 |
| 2rln | 69.16 | 48.95 |
| 4hyb | 91.15 | 96.25 |
| 1uef | 91.20 | 90.19 |
| 1rbd | 93.45 | 79.66 |
| 3fqx | 92.04 | 92.50 |
| 3kmz | 93.53 | 97.50 |
| 3cmh | 71.07 | 72.50 |
| 3ejh | 85.14 | 82.34 |
| 2vpg | 66.76 | 76.44 |
| 3l6f | 95.19 | 98.50 |
| 3bgm | 93.17 | 93.62 |
| 1z3p | 94.53 | 94.56 |
| 3f9y | 85.97 | 95.00 |

### Section S6. The result of relaxation

**Table S9.** Summary statistics of structure quality for all unnatural residues in the independent cyclic peptide test set.

| Residue | PDBID | Clashscore[a] | Poor rotamers | Favored rotamers | Ramachandran outliers | Ramachandran favored | MolProbity[b] |
|---|---|---|---|---|---|---|---|
| ABA | 2m2h | 52.63 / 0 | 0 / 0 | 12 / 11 | 0 / 0 | 4 / 4 | 2.2 / 0.5 |
| DLE | 2m6d | 165.29 / 32.52 | 2 / 1 | 4 / 4 | 1 / 0 | 1 / 3 | 4.88 / 2.92 |
| DPR | 6dze | 86.81 / 3.42 | 3 / 1 | 10 / 11 | 0 / 0 | 11 / 11 | 3.42 / 1.78 |
| HYP | 1k64 | 11.11 / 0 | 2 / 0 | 13 / 15 | 1 / 0 | 12 / 12 | 2.04 / 1.79 |
| NH2 | 1omc | 51.15 / 2.55 | 1 / 0 | 22 / 23 | 1 / 0 | 13 / 14 | 3.39 / 1.65 |
| ORN | 2pm5 | 114.35 / 4.49 | 0 / 0 | 21 / 21 | 0 / 0 | 21 / 21 | 2.52 / 1.23 |
| PCA | 1big | 32.14 / 1.78 | 3 / 0 | 29 / 31 | 0 / 0 | 32 / 32 | 3.12 / 1.33 |

[a] The values before relaxation are listed first, followed by the values after relaxation. This format applies to all subsequent columns in the table.

[b] The MolProbity score integrates clash scores, rotamer evaluations, and Ramachandran assessments, effectively reflecting a structural model's overall quality and reliability.

**Table S10.** The bond lengths of the residues' covalent bonds.

| Residue | PDB ID | Atom pair[a] | Bond(Crystal)[b] | Bond(Unrelax / Relaxed)[c] |
|---|---|---|---|---|
| ABA | 2zol | CB-CG | 1.52 | 1.54 / 1.53 |
| ALC | 4hyb | CB-CG | 1.53 | 1.52 / 1.55 |
| ALY | 4lka | NZ-CH | 1.35 | 1.30 / 1.36 |
| CSO | 3kmz | SG-OD | 1.79 | 1.68 / 1.79 |
| MLZ | 3kmt | NZ-CM | 1.47 | 1.44 / 1.47 |
| MSE | 2oq1 | CE-SE | 1.90 | 2.01 / 1.84 |
| NVA | 1z3p | CG-CD | 1.51 | 1.51 / 1.51 |
| ORN | 3idj | CB-CG | 1.54 | 1.55 / 1.55 |
| PRK | 3pdh | NZ-CAL | 1.33 | 1.34 / 1.38 |
| PTR | 2ci9 | OH-P | 1.62 | 1.59 / 1.59 |

| | | | | |
|---|---|---|---|---|
| SEP | 3fqu | OG-P | 1.61 | 1.64 / 1.61 |
| TPO | 2k7l | OG1-P | 1.61 | 1.62 / 1.61 |

[a] In unnatural amino acid residues, the atom pairs that form covalent bonds with the natural amino acids through modification groups. The bond is in Å.

[b] The bond lengths of the Atom pairs in the crystal structure.

[c] The bond lengths of the Atom pairs before and after relaxation.

**Table S11.** All RMSD for the relaxed structure in the independent cyclic peptide test set.

| PDB ID | $RMSD_{C\alpha}$ | $RMSD_{all\text{-}atom}$ | $RMSD_{unAA}$ |
|---|---|---|---|
| 2ew4 | 1.075 | 2.059 | 1.866 |
| 2msq | 1.890 | 2.673 | 2.059 |
| 6dz9 | 2.288 | 4.648 | 3.879 |
| 1t9e | 2.076 | 2.992 | 2.529 |
| 2m2s | 2.224 | 3.408 | 2.224 |
| 2pm5 | 0.890 | 1.531 | 1.613 |
| 4e86 | 0.818 | 1.711 | 1.161 |
| 1big | 1.196 | 1.990 | 1.729 |
| 3wng | 1.179 | 1.696 | 3.323 |
| 2m2x | 3.774 | 6.722 | 5.163 |
| 5ug3 | 3.362 | 4.885 | 3.438 |
| 4e83 | 0.913 | 1.581 | 0.651 |
| 2m6d | 1.870 | 2.832 | 2.841 |
| 2crd | 1.112 | 2.190 | 0.923 |
| 1p9g | 1.180 | 2.031 | 4.762 |
| 2mg6 | 1.643 | 2.365 | 3.747 |
| 1omc | 1.022 | 2.568 | 1.566 |
| 2m2g | 0.645 | 1.881 | 1.401 |
| 2n8e | 2.216 | 3.248 | 6.410 |

| | | | |
|---|---|---|---|
| 2j15 | 1.594 | 3.114 | 1.746 |
| 6dze | 2.941 | 4.624 | 4.399 |
| 6dza | 3.961 | 5.808 | 4.857 |
| 2m7j | 3.724 | 6.372 | 4.142 |
| 2m2h | 0.758 | 2.398 | 1.618 |
| 5n99 | 7.940 | 7.947 | 8.342 |
| 6dzc | 3.028 | 5.425 | 4.014 |
| 2m1p | 0.658 | 2.103 | 1.506 |
| 2m62 | 1.892 | 2.880 | 1.085 |
| 2mfx | 3.667 | 4.170 | 4.412 |
| 2ih6 | 2.354 | 3.742 | 3.684 |
| 6my2 | 1.593 | 2.839 | 4.121 |
| 1k64 | 5.621 | 6.808 | 10.318 |
| 6dzb | 2.272 | 4.522 | 1.840 |
| 3lo6 | 0.635 | 1.236 | 0.452 |

**Table S12.** All RMSD for the relaxed structure in the linear peptide test set.

| PDB ID | $RMSD_{C\alpha}$ | $RMSD_{all\text{-}atom}$ | $RMSD_{unAA}$ |
|---|---|---|---|
| 6cmh | 5.704 | 6.222 | 3.221 |
| 2k7l | 3.404 | 4.461 | 5.276 |
| 2fx8 | 0.928 | 1.581 | 1.026 |
| 1fev | 0.400 | 1.526 | 0.422 |
| 1jy9 | 2.388 | 2.682 | 1.819 |
| 2zok | 1.158 | 2.136 | 0.451 |
| 3avr | 1.016 | 1.466 | 1.304 |
| 3pdh | 3.021 | 3.926 | 0.460 |
| 1mxq | 0.995 | 1.735 | 0.970 |
| 2h6n | 5.095 | 8.096 | 10.823 |
| 2ci9 | 0.852 | 1.660 | 1.332 |
| 3kmt | 0.732 | 1.031 | 0.996 |
| 3u5n | 0.867 | 1.833 | 2.841 |
| 2zsw | 1.947 | 3.754 | 1.273 |
| 3nti | 9.000 | 11.259 | 5.656 |
| 1v50 | 2.058 | 3.039 | 2.996 |
| 4lka | 0.819 | 1.193 | 2.046 |
| 3fqu | 0.842 | 1.913 | 3.442 |
| 3idj | 4.634 | 4.853 | 7.876 |
| 2zol | 0.460 | 0.611 | 0.661 |

| | | | |
|---|---|---|---|
| 2oq1 | 1.351 | 2.000 | 2.086 |
| 4jaa | 2.101 | 2.290 | 2.872 |
| 2nor | 0.467 | 1.873 | 3.018 |
| 2rln | 2.074 | 3.755 | 6.451 |
| 4hyb | 0.387 | 0.986 | 1.563 |
| 1uef | 0.654 | 1.693 | 1.798 |
| 1rbd | 0.602 | 1.398 | 1.231 |
| 3fqx | 0.515 | 1.314 | 2.362 |
| 3kmz | 1.001 | 3.200 | 1.781 |
| 3cmh | 3.404 | 4.184 | 2.616 |
| 3ejh | 9.351 | 10.515 | 10.592 |
| 2vpg | 4.404 | 4.952 | 3.550 |
| 3l6f | 0.702 | 1.271 | 1.215 |
| 3bgm | 0.783 | 1.559 | 1.525 |
| 1z3p | 0.491 | 1.553 | 1.637 |
| 3f9y | 0.412 | 1.785 | 0.597 |
